# Engineering compact *Physalis peruviana* (goldenberry) to promote its potential as a global crop

**DOI:** 10.1101/2025.08.15.670557

**Authors:** Miguel Santo Domingo, Blaine Fitzgerald, Gina M. Robitaille, Srividya Ramakrishnan, Kerry Swartwood, Nicholas G. Karavolias, Michael C. Schatz, Joyce Van Eck, Zachary B. Lippman

## Abstract

EN - Goldenberry (*Physalis peruviana*) produces sweet, nutritionally-rich berries, yet like many minor crops, is cultivated in limited geographical regions and has not been a focus of breeding programs for trait enhancement. Leveraging knowledge of plant architecture-related traits from related species, we used CRISPR/Cas9-mediated gene editing to generate a compact ideotype to advance future breeding efforts and agricultural production. Goldenberry growers will benefit from these compact versions because it optimizes per plot yield, facilitating larger-scale production to meet rising consumer popularity and demand.

SP - La uchuva (*Physalis peruviana*) produce frutos dulces y ricos en nutrientes, pero, igual que muchos cultivos minoritarios, se cultiva en zonas geográficas limitadas y no ha sufrido un proceso de mejora. Aprovechando conocimientos sobre rasgos relacionados con la arquitectura vegetal de especies relacionadas, hemos usado edición génica mediante CRISPR/Cas9 para generar un ideotipo compacto para promover futuros esfuerzos en su mejora y en producción agrícola. Los productores de uchuva se podrán beneficiar de estas versiones compactas ya que optimiza el rendimiento por parcela, facilitando así la producción a una mayor escala para cubrir la creciente popularidad y demanda de los consumidores.

## INTRODUCTION

A limited number of species dominate global crop production (FAO, 2024) resulting in a fragile food supply. However, minor crops, sometimes referred to as indigenous, local, or orphan crops, are relied upon in regional contexts, where, although not fully domesticated, they provide nutritional dietary diversity. Some minor crops, such as quinoa and dragon fruit, have become popular with worldwide consumers because of increased interest in new flavors, textures, and their health benefits.

Goldenberry (*Physalis peruviana*), a member of the crop-rich Solanaceae family, is a minor crop indigenous to South America that is often touted as a superfood because of its high nutritional value (Shenstone, Lippman and Van Eck, 2020). It produces round, smooth, yellow-orange colored berries that are surrounded by a husk (inflated calyx), similar to other species in the *Physalis* genus, and its fruit flavor is reminiscent of other tropical fruits such as pineapple and mango. Depending on the region where it is grown, goldenberry is known by locally recognized names such as uchuva (Colombia) and Cape gooseberry (South Africa). While goldenberries have been historically grown in the Andean region near its evolutionary origin, they have undergone little domestication, as evidenced by its large, wild, unmanageable growth habit that complicates large-scale production (Flores Roncancio et al., 2020). Currently, Colombia is the main producer of goldenberries, producing more than 20,000 tons of fresh fruit in 2022 with 40% exported (Resilience BV, 2024); however, additional production occurs in Peru, Ecuador, South Africa and India. Improvement of its growth habit to make it more manageable in large-scale production would have a positive impact on dietary diversification with greater availability of this highly nutritious fruit.

Modification of traits in many plant species, including minor and major solanaceous crops, has been readily achievable by the application of genome editing technologies (Lemmon et al., 2018; Kwon et al., 2020). For example, we showed that CRISPR-engineered mutations of the classical stem length regulator *ERECTA* (Torii et al., 1996) in groundcherry (*Physalis grisea*) and tomato (*Solanum lycopersicum*) resulted in a more compact (determinate) growth habit (Kwon et al., 2020). Based on this outcome, engineering mutations of the two *ERECTA* genes in tetraploid goldenberry could also result in plants with a modified, and more agriculturally desirable, plant architecture.

We report here the development of high-quality genome assemblies alongside transformation and genome-editing approaches, which we used to rapidly modify goldenberry plant architecture by targeting the two sub-genome copies of *ERECTA*. To give breeders and growers a head-start with our improved, compact goldenberry without any regulatory delays, we have already acquired clearance by the US Department of Agriculture (USDA) to document that our varieties are not plant pests and no plant pests remain integrated into the plant genome, so that they are not regulated under 7 CFR Part 340. In addition, we also seek approval by the US Food and Drug Administration (FDA) to enable growers to immediately move forward with commercial production promoting rapid adoption and new market opportunities.

## ECONOMIC AND BREEDING POTENTIAL OF GOLDENBERRY

Goldenberry fruits are small (4 to 10 grams) (Flores Roncancio et al., 2020) and round with a bright yellow-orange color, and have a unique flavor and nutritional profile (**Figure 1A**) (Flores Roncancio et al., 2020; Shenstone, Lippman and Van Eck, 2020). Their appeal as a new berry for consumers worldwide has expanded their accessibility to markets and grocery stores beyond South America. In particular, in the United States, goldenberries are now sold year round (**Figure 1B**), revealing its high economic potential.

**Figure 1.**
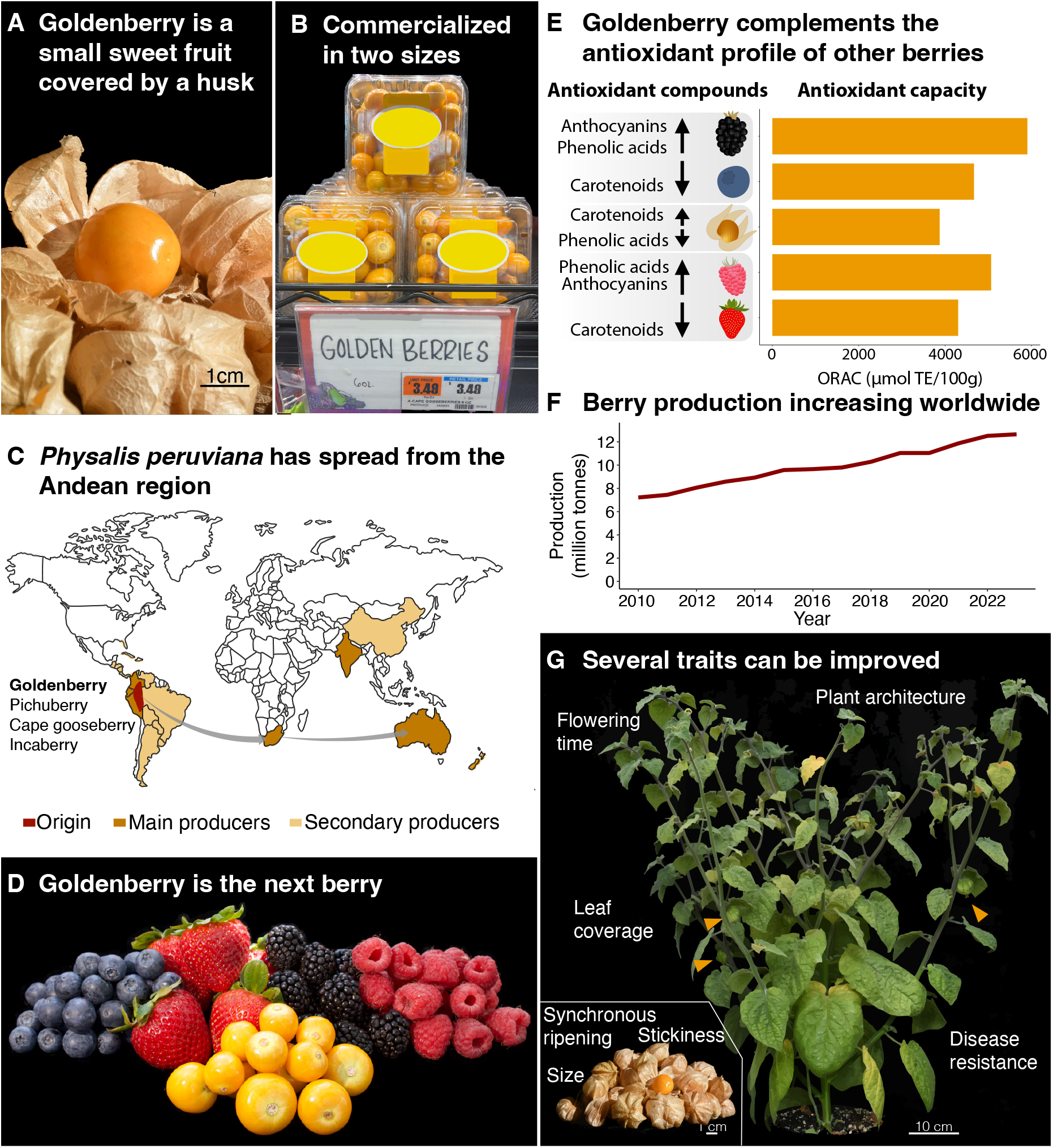
Goldenberry (*Physalis peruviana*): an emerging commercial berry crop and a promising candidate crop to help globalize using genome editing. **A)** Goldenberry fruits covered by a papery husk, typical of the inflated calyx syndrome (ICS) in the genus *Physalis* and other Solanaceae species. A fruit with its husk removed is also shown. **B)** Commercial goldenberries are assorted in two sizes: “small” and “large” types (Supplementary Figure S3). **C)** Historical origin, distribution, and producers of goldenberry. **D)** Commercial goldenberry is comparable and complementary to other berries in terms of fruit size and consumer interest. **E)** Oxygen Radical Absorbance Capacity (ORAC) measurements of antioxidants of the four major berries compared to goldenberry. Arrows represent the comparative contribution to antioxidant capacity of the different compounds in the different groups of berries. **F)** Growing worldwide production of berries (strawberries, raspberries, blueberries) from 2010 to 2023. **G)** Several plant and fruit traits which could be improved in goldenberry. Orange arrows point to goldenberry fruits.

Native to Peru, goldenberry grows wild between 1,500 and 3,000 m in elevation in the Andean countries of Colombia and Peru, where it has been consumed for at least several centuries dating to the Inca Empire (Legge, 1974). After European colonization in the 18th century, goldenberry was brought to South Africa and other parts of the world including India and Australia (**Figure 1C**). Its small, round fruits, are highly nutritious with health beneficial levels of antioxidants, carotenoids, and micronutrients (Ezbach et al., 2018; Haytowitz and Bhagwat, 2010; Gündeşli et al., 2019), which puts goldenberry in the superfood category, together with other commercial berries (**Figure 1D, E**). Similar to other berries, goldenberry is primarily consumed as fresh fruit, however, they are also used to make jams, condiments such as chutney, and are also available as dried fruit (raisins) (Van Eck, 2022). The global growing appeal of berries is evidenced by their expanding production and availability in markets (**Figure 1F**) (FAOSTAT, 2024), positioning goldenberries to become a highly sought after crop by farmers and distributors. However, for its economic value and large-scale agricultural production to be realized, critical trait enhancements (e.g. synchronous ripening, reduction of sticky acylsugars on fruits, elimination of fruit cracking, and a modified plant architecture) are required through plant breeding, including new technologies such as genome editing (**Figure 1G**).

## ENGINEERING A COMPACT GOLDENBERRY GROWTH HABIT

To improve its growth habit, we focused on engineering mutations in the goldenberry *ERECTA* gene (*Pper-ER*). We based this approach on previous work where we targeted *ERECTA* in sister species tomato and groundcherry to modify plant architecture (Kwon et al., 2020). However, goldenberry presented several challenges. First, as an allotetraploid (4x = 2n = 48) (Sanchez-Betancourt and Núñez Zarantes, 2022), the possibility of two divergent copies of *Pper-ER* (one for each subgenome) in goldenberry makes editing more challenging, as both gene copies might require independent sets of guide RNA (gRNA), coupled with more complex downstream genetic fixation of desired alleles compared to a diploid. Second, there was the need to establish efficient plant regeneration and transformation methods for this species along with generating a high quality reference genome.

We generated high-quality genomes for two ecotypes, *India* and *South Africa*. Using long PacBio HiFi reads, we estimated the haploid genome size to be ∼1.34 Gbp. Assemblies yielded ∼5.3 Gbp genomes, representing the four distinct haplotypes, with an average contig N50 of ∼48.0 Mbp and the longest contigs approaching complete chromosomes or chromosome arms (max: 226.54 Mbp) (**Figure 2A**) (**Supplementary Table S1**). Genome completeness analysis showed ∼98.3% completeness, with 94.1% of genes duplicated, consistent with tetraploidy. The haplotype resolved assemblies and their corresponding gene duplications enabled us to study haplotypic variation in the genes (**Supplementary Table S2**). Importantly, we detected two highly similar *Pper-ER* genes, *Pper-ER-A* and *Pper-ER-B*, allowing the design of a four-gRNA construct for simultaneous targeting (**Supplementary Table S3**). In parallel, we developed plant regeneration, transformation and gene-editing approaches. Regeneration efficiency was similar in both ecotypes (36% and 37% of explants regenerating at least one shoot in *South Africa* and *India*, respectively), but editing was more efficient in the *India* ecotype. We did not detect edits in *South Africa ERECTA* genes, but we recovered, among others, two null alleles in *Pper-ER* in the *India* ecotype: one base pair deletion in *Pper-er-A* and one base pair insertion in *Pper-er-B*, in the first and third exons, respectively (**Figure 2B**).

**Figure 2.**
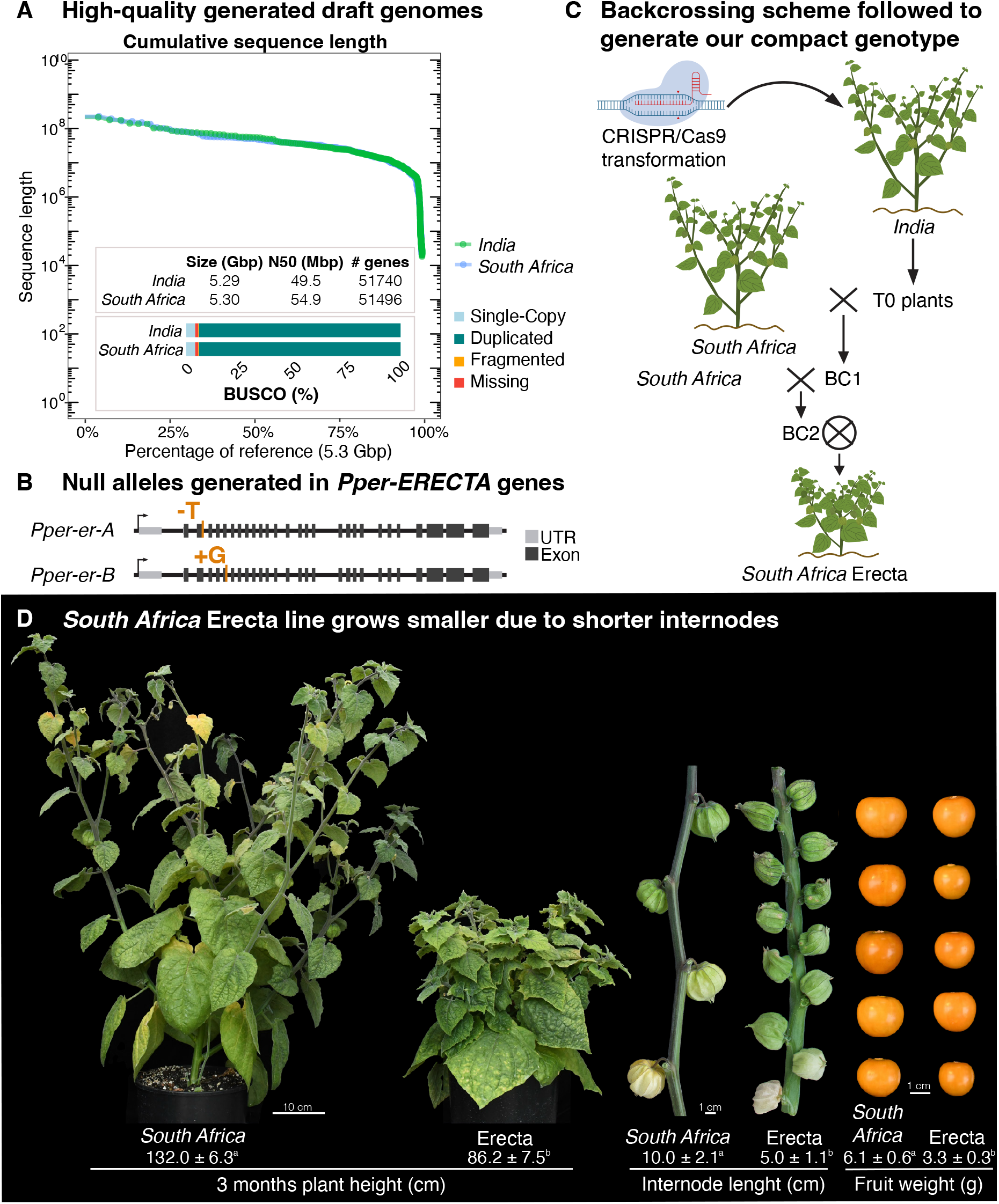
Establishment of reference genomes, genome editing, and a compact “Erecta” line to improve goldenberry cultivation. **A)** High-quality genome assemblies of the tetraploid parental lines *South African* and *India*, featuring maximum contig lengths approaching chromosome arm lengths and high BUSCO completeness, indicating assembly integrity and comprehensive gene content. **B)** Crossing scheme followed to introduce the CRISPR/Cas9-engineered *erecta* mutant alleles into the commercial *South Africa* ecotype to develop two Erecta varieties. **C)** Generated null mutant alleles in both *Pper-ERECTA* genes. **D)** Phenotypes of the *South Africa* parental line compared to its corresponding Erecta line, showing a drastic reduction in plant size and internode length. Fruits also become smaller, but are within the range of current commercial types. Plant height: n = 20. Internode length: n = 50. Fruit weight: n = 50. Small letters designate significantly different groups. Significant groups were defined as p-value < 0.05.

We then introgressed both null alleles into the *South Africa* line since there is some preference for their fruit flavor. After two rounds of backcrossing the edited lines to *South Africa* and a final self-pollination, we recovered plants with substantially increased compactness, which we named “Erecta” to denote this genotype as a new line (**Figure 2C**). In subsequent generations, we made selections from the compact plant lines based on fruit flavor, resulting in stable, compact varieties of *India-* and *South Africa*-like flavored fruits. Finally, we sequenced the genome of the *South Africa* Erecta line to demonstrate absence of the transgene (T-DNA insertion) and as a non-transgenic resource for future development through plant breeding (**Supplementary Table S1, Supplementary Table S2, Supplementary Figure S1**).

Three-month-old *South Africa* Erecta plants were 35% shorter than *South Africa* parental line plants (86.2 +-7.5 cm and 132.0 +-6.3 cm, respectively) (**Figure 2D, Supplementary Figure S2**). This decrease resulted from a 50% reduction in internode length compared to the parental line (5.0 +-1.1 cm in Erecta 10.0 += 2.1 cm in *South Africa*), consistent with the effect of *er* mutations in other species (Kwon et al., 2020). Importantly, the fruit fecundity (total amount of fruits produced per plant) was not affected by the mutations (**Supplementary Figure S2D**). However, similar to other *erecta*-edited crops, fruit size was also reduced proportionally; *South Africa* Erecta fruit weighed ∼50% of *South Africa* parental line fruit (3.3 +-0.3 g and 6.1 +-0.6 g, respectively).

Despite the reduced fruit size, the goldenberry Erecta lines are still valuable for breeding and production. Goldenberries currently on the market are sold in two sizes, which we designated as “large” and “small” types (**Figure 1B**). The average fruit weight of the large type is ∼7.4 g, while the small type is ∼3.8 g (**Supplementary Figure S3**). Our Erecta plants produced, on average, 3.3 g fruits (**Figure 2D, Supplementary Figure S3**), only slightly less than those commercially available, making them well-suited for immediate commercial evaluation. Thus, the *South Africa* Erecta germplasm will provide a foundation for subsequent improvement of this flavorful and highly nutritious fruit.

In summary, using CRISPR-Cas9-mediated gene editing of the *India* genotype followed by several cycles of introgression and selection into the *South Africa* genotype because of its preferred fruit flavor, we have generated two stably modified compact germplasm with fruits similar to commercially available ones. With this new Erecta germplasm, planting density can be increased, while also avoiding the need for laborious staking and trellising of large, bushy plants (**Figure 3**).

**Figure 3.**
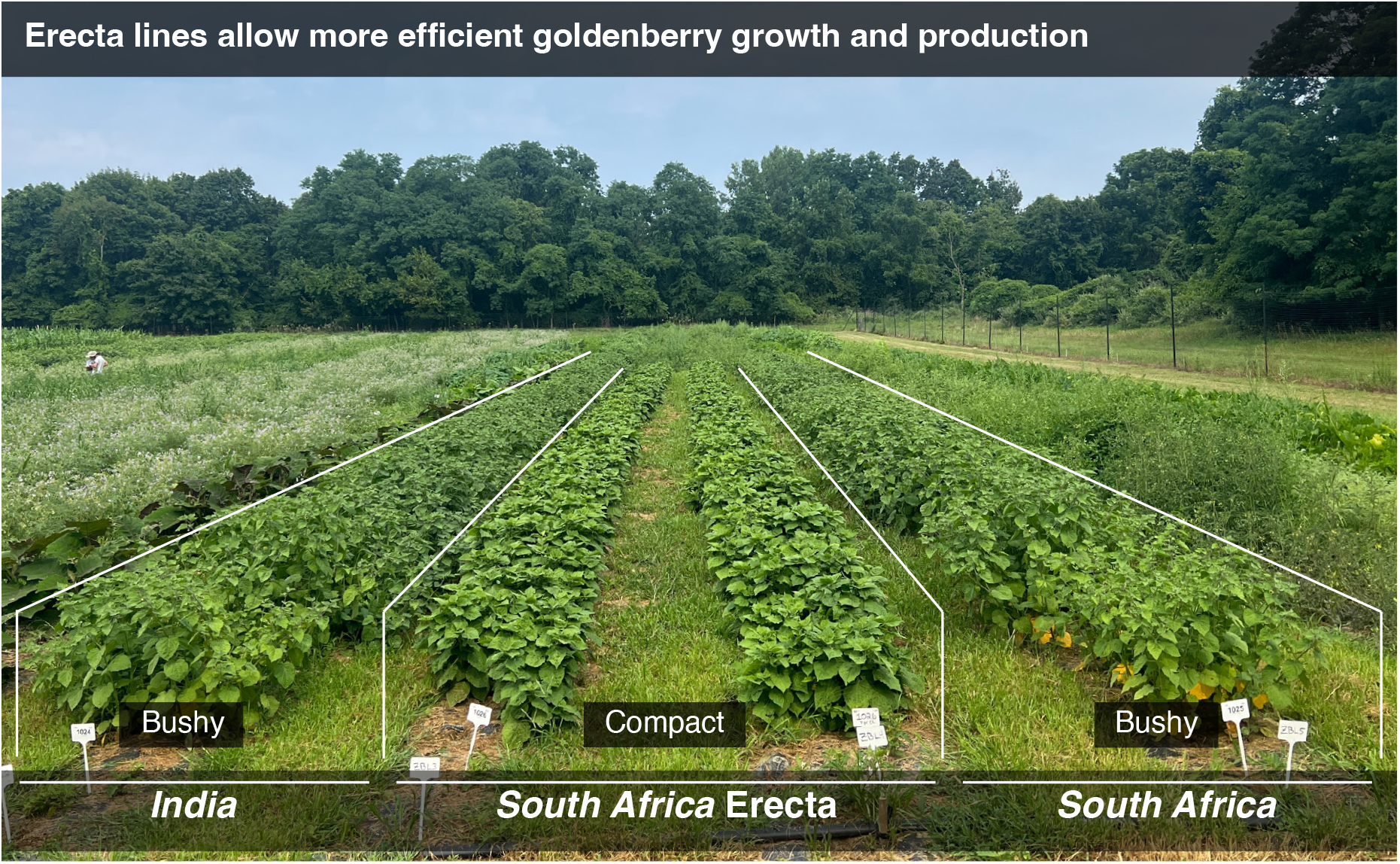
Demonstration of field-based row production of the new *South Africa* Erecta line (two middle rows) compared to the much larger and bushier *South Africa* (right row) and *India* (left row) parental lines. Note the shorter, prostrate growth habit of the *South Africa* Erecta line, facilitating high-density planting, plant maintenance, and harvestability.

## IMPACT

We mutated *Pper-ER* and established a pre-breeding program for compact goldenberry plants by integrating information from related major and minor solanaceous crop species, and following an introgression scheme. These new Erecta lines can be used as a starting point to develop new varieties through traditional breeding and possibly additional gene editing to improve other traits, or directly in the production of goldenberries.

For further improvement, fruit size is the most obvious next target. As seen in *P. grisea* (Lemmon et al., 2018), the targeting of *PperCLV1* could help restore and potentially increase fruit size in the Erecta lines. Additionally, other genes responsible for tomato fruit characteristics could further customize fruit size and shape (Mauxion, Chevalier and Gonzalez, 2021). Additional improvements include the modification of acylsugar metabolism to reduce fruit stickiness or accelerated flowering for earlier harvests (**Figure 1G**).

Apart from goldenberry, several other minor crops could benefit from the presented scheme for improvement of undesirable traits. Already, genomes and transformation approaches have been developed (or are under development) for a range of minor crops, such as passion fruit (Xia et al., 2021; Manders et al., 1994), fonio (Abrouk et al., 2020; Ntui et al., 2017), and groundcherry (Dale et al. 2024) leveraging the potential of gene editing to expand the benefits of local (agri)culture worldwide.

## AUTHOR CONTRIBUTIONS

MCS, JVE, and ZBL conceived the project and designed the experiments. SR and MCS generated the genomes. KS and JVE developed the transformation methods and generated the edited plants. BF, GMR, and ZBL developed the Erecta genotypes. MSD and BF collected phenotypic data. MSD performed the data analysis and wrote the original draft. All the authors revised and edited the manuscript.

## ACKNOWLEDGEMENTS

We thank members of the Lippman laboratory for discussions; B. Seman for technical support; T. Mulligan, K. Schlecht, and S. Qiao for assistance with plant care. We thank G. Vogel of Cornell University for guidance on terminology related to the edited lines we developed. This work was supported by the National Science Foundation Plant Genome Research Program grant IOS-2216612 to J.V.E., M.C.S. and Z.B.L. We thank the Indigenous peoples of South America on whose ancestral lands P. peruviana grow.

## CONFLICT OF INTEREST

All authors declare no competing interests.

## Notes

### Competing Interest Statement

The authors have declared no competing interest.

### Summary of Updates

More information has been added about the methodology.

