## Supplementary Figure for "Engineering compact *Physalis peruviana* (goldenberry) to promote its potential as a global crop"

### SUPPLEMENTARY FIGURES

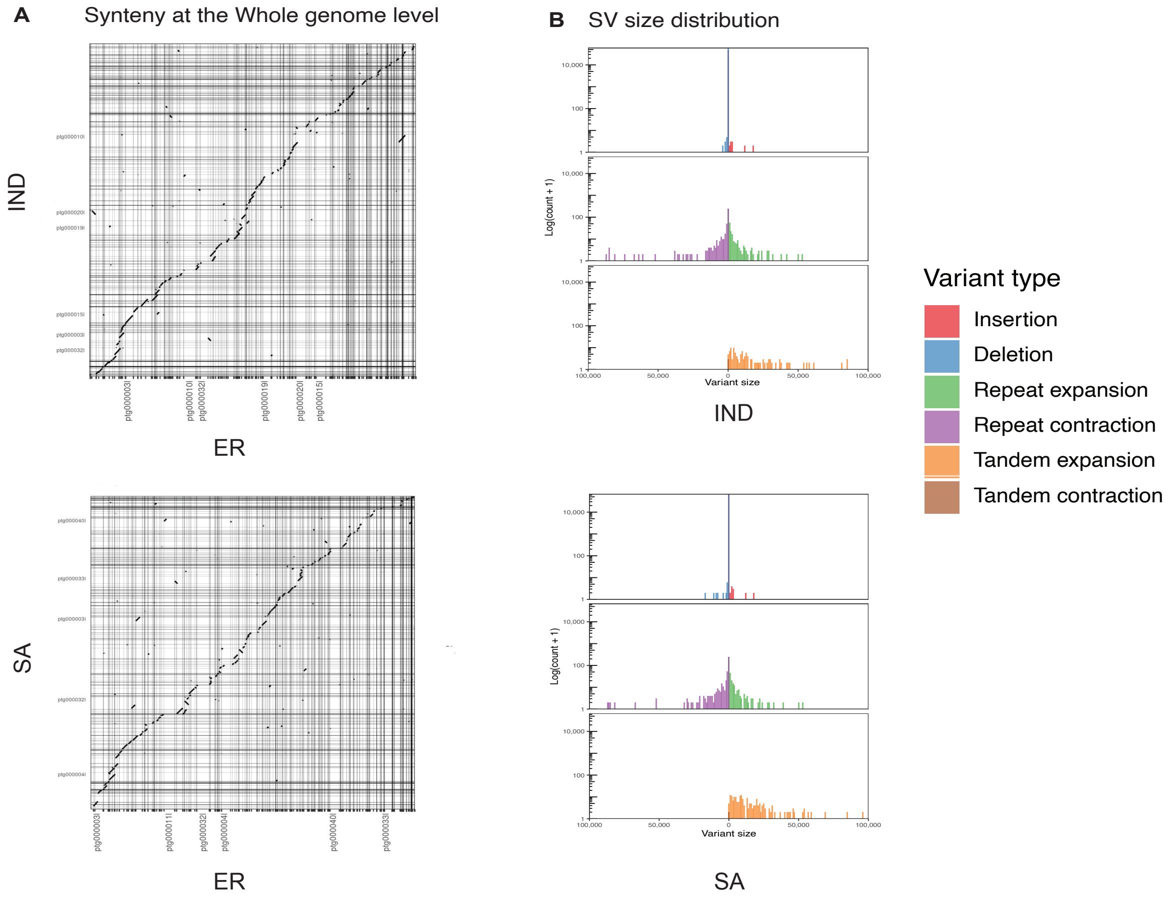


**Supplementary Figure S1. Genotype comparison from whole genome sequencing of the parental ecotypes *India* and *South Africa* and the *South Africa* Erecta line. Minimal structural variation (SV) is found between the lines**. **A**) Whole-genome synteny analysis of the *India* and *South Africa* ecotypes compared to the *South Africa* Erecta line. **B**) SV summary of the parental *India* and *South Africa* ecotypes as compared to the *South Africa* Erecta line.

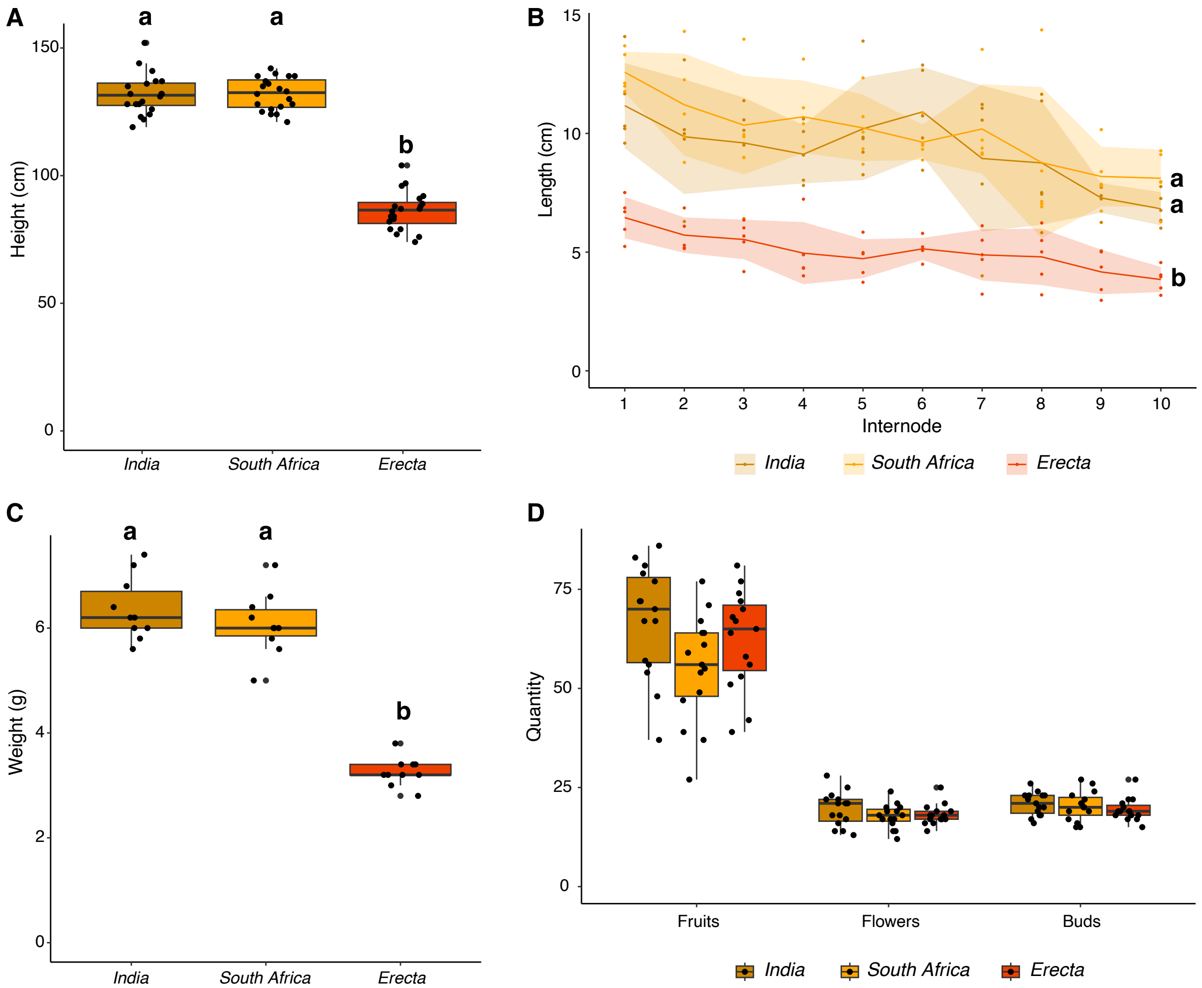


**Supplementary Figure S2. Phenotypic comparison of the *South Africa* Erecta line with *South Africa* and *India* parental lines**. **A**) Plant height (cm) **B**) Internode length (cm) **C**) Fruit weight (g) **D**) Number of fruits, flowers and buds, as a measure of fecundity. All phenotypes were measured in 90-day old plants (60 days after transplant to the field). Small letters designate significantly different groups (p-value < 0.05).

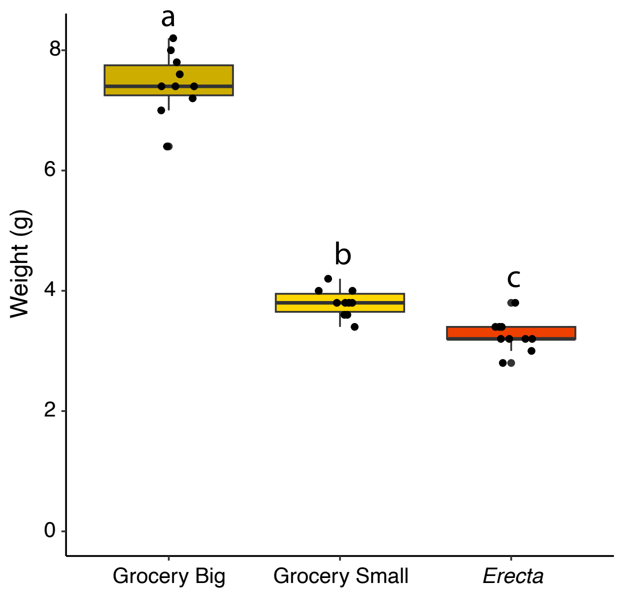


**Supplementary Figure S3. Fruit weight (g) comparison of the *South Africa* Erecta line with the two size types of currently available commercial goldenberries**. Small letters designate significantly different groups (p-value < 0.05).
