## Supplementary material for "Engineering compact *Physalis peruviana* (goldenberry) to promote its potential as a global crop": Materials and Methods

### Plant material and growing conditions.

Seeds were sown in soil in 96-cell plastic flats and grown to 4-week-old seedlings in greenhouses. The seedlings were then transplanted directly to the field at Cold Spring Harbor Laboratory, New York. Greenhouse conditions were long-day (16 h light, 26–28 °C followed by 8 h dark, 18–20 °C; 40–60% relative humidity) with natural light supplemented with artificial light from high-pressure sodium bulbs (~250 µmol m−2 s−1). Plants in the field were grown under drip irrigation and standard fertilizer regimens, and were used for quantifications of internode length, plant height, fecundity, and fruit weight.

### Generation of CRISPR/Cas9 induced mutations

CRISPR Guide RNAs to target *ERECTA* in *P. peruviana* were designed using Geneious. The four-guide construct was cloned following the Golden Gate cloning approach as described in Brooks et al, 2014.

For *Physalis peruviana* transformation, seeds were surface sterilized and germinated in vitro according to methods previously reported for *Physalis pruinosa* (Starwood and Van Eck, 2019). One day (preculture) prior to transformation, hypocotyls from 10 – 11-day-old seedlings (before emergence of first true leaves) were sectioned into 0.5 cm lengths and transferred to Plant Regeneration Medium 1 (PM1) that contained the following components per liter: 4.3 g MS salts, 100 mg myo-inositol, 1 ml of a 1000X solution of modified Nitsch and Nitsch vitamins (per 100 ml: glycine 0.2 g, nicotinic acid 1 g, pyridoxine–HCl 0.05 g, thiamine-HCl 0.05 g, folic acid 0.05 g, d-biotin 0.004 g, pH 7.0), 20 g sucrose, 2 mg benzylaminopurine (BA, Phytotech), 0.5 naphthaleneacetic acid (NAA, Phytotech) and 8 g TC Agar (Phytotech).

In parallel, CRISPR-Cas9 constructs with four guides were introduced by electroporation into electrocompetent *Agrobacterium tumefaciens* AGL1 (Lazo, Stein and Ludwig 1991) and prepared for transformation as previously reported (Starwood and Van Eck, 2019).

For transformation, the 1-day precultured explants were incubated in the *Agrobacterium*/2% MSO suspension for 5 min then transferred to a sterile paper towel to briefly drain excess suspension. The explants were placed back onto PM1 and cultured in the dark at 19 °C for 48 h. Plates were sealed with parafilm for the cocultivation period. After 48 h, the explants were transferred to selective PM1 that contained 300 mg/l timentin and 200mg/l kanamycin, and designated PM1K as kanamycin is used for the selection of transformants. One month later, they were cultured on PM2K medium, which is similar to PM1K with no NAA included.

Cultures were transferred to freshly prepared PM1K and PM2K every two weeks in either Petri plates or Magenta boxes depending upon size of shoots regenerating from the explants.

When regenerated shoots were approximately 2 cm tall, they were excised from explants and transferred to selective rooting medium that contained the following components per liter: 4.3 g MS salts, 1 ml of a modified Nitsch and Nitsch vitamins solution (see above), 30 g sucrose, 300 mg timentin, 200 mg kanamycin, 8 g Difco Bacto agar (Becton, Dickinson and Company, Franklin Lakes, NJ). Magenta boxes that contained 62.5 mL of medium were used for the rooting phase.

Once rooted, for acclimation to greenhouse conditions, culture medium was washed before transfer to moistened soilless mix. The plants were immediately covered with plastic domes, which were gradually removed over the course of 5 days. Plants were maintained in a growth chamber for 2 – 3 weeks then transferred to a greenhouse.

Unless otherwise noted, the following conditions were followed. The pH of all media was adjusted to 6.0 before autoclaving. For all media, carbenicillin, BA, NAA, kanamycin, and timentin were dispensed from filter sterilized stock solutions into autoclaved medium cooled to 55 °C. The size of the Petri plates used was 100 mm × 20 mm. Plates were sealed with Micropore tape (Amazon). All cultures were maintained at 24 ± 2 °C under a 16 h light/8 h dark photoperiod at 57–65 uE m−2 s−1.

**DNA extraction**

For extraction of high-molecular-mass DNA, young leaves were collected from 21-day-old light-grown seedlings. Before tissue collection, seedlings were etiolated in complete darkness for 48 h. Flash-frozen plant tissue was ground using a mortar and pestle and extracted in four volumes of ice-cold extraction buffer 1 (0.4 M sucrose, 10 mM Tris-HCl pH 8, 10 mM MgCl2 and 5 mM 2-mercaptoethanol). Extracts were briefly vortexed, followed by dounce homogenization on ice and filtered twice through a single layer of Miracloth (Millipore Sigma). Filtrates were centrifuged at

4,000 rpm for 20 min at 4 °C, and pellets were gently resuspended in 1 ml of extraction buffer 2 (0.25 M sucrose, 10 mM Tris-HCl pH 8, 10 mM MgCl2, 1% Triton X-100, and 5 mM 2-mercaptoethanol). Crude nuclear pellets were collected by centrifugation at 12,000*g* for 10 min at 4 °C and washed by resuspension in 1 ml of extraction buffer 2 followed by centrifugation at 12,000*g* for 10 min at 4 °C. Nuclear pellets were resuspended in 500 ml of extraction buffer 3 (1.7 M sucrose, 10 mM Tris-HCl pH 8, 0.15% Triton X-100, 2 mM MgCl2 and 5 mM 2-mercaptoethanol), layered over 500 ml extraction buffer 3 and centrifuged for 30 min at 16,000*g* at 4 °C. The nuclei were resuspended in 2.5 ml of nuclei lysis buffer (0.2 M Tris pH 7.5, 2 M NaCl, 50 mM EDTA and 55 mM CTAB) and 1 ml of 5% Sarkosyl solution and incubated at 60 °C for 30 min.

To extract DNA, nuclear extracts were gently mixed with 8.5 ml of chloroform:isoamyl alcohol solution (24:1) and slowly rotated for 15 min. After centrifugation at 4,000 rpm for 20 min, 3 ml of aqueous phase was transferred to new tubes and mixed with 300 ml of 3 M NaOAc and 6.6 ml of ice-cold ethanol. Precipitated DNA strands were transferred to new 1.5 ml tubes and washed twice with ice-cold 80% ethanol. Dried DNA strands were dissolved in 100 ul of elution buffer (10 mM Tris-HCl, pH 8.5) overnight at 4 °C. The quality, quantity and molecular mass of DNA samples were assessed using Nanodrop (Thermo Fisher Scientific), Qubit (Thermo Fisher Scientific) and pulsed-field gel electrophoresis (CHEF Mapper XA System, Bio-Rad) according to the manufacturer's instructions.

**Genome assembly and annotation**

Reference draft assemblies for *Physalis peruviana India ecotype*, *South Africa* ecotype and *South Africa* Erecta line were generated using long-read sequencing (Pacific Biosciences, USA).

Genome size, ploidy, and heterozygosity of *P. peruviana* were estimated using GenomeScope 2.0 (Ranallo-Benavidez et al., 2020) with 21 bp kmer profiles from PacBio HiFi reads and ploidy set to 4. Assemblies were generated using *hifiasm* (Cheng et al., 2021) with options -l=3 and --n-hap=4, followed by screening to remove bacterial, fungal, mitochondrial, and chloroplast contigs. Genome completeness was evaluated with BUSCO v5 (Manni et al., 2021) using the Solanales_odb10 dataset. Structural variants were identified by aligning *India* and *South Africa* genome assemblies to the *South Africa* Erecta reference with Minimap 2 (Li, 2018) and calling variants using paftools and Assemblytics (Nattestad and Schatz, 2016) respectively.

For genome annotation, orthologs with coverage above 50% and 75% identity were lifted from *Physalis grisea* genome (He et al., 2022) LiftON (Chao et al., 2025) and refined using protein and gene microsynteny support. The completeness of the gene models was determined by assessing single-copy orthologs using BUSCO5 (Manni et al., 2021).

**Phenotyping**

Phenotyping was performed in 90 to 100-day old plants.

To quantify internode length, ten internodes above the first true leaf were measured for five plant replicates from each of the three genotypes. Distances between each adjacent pair of petioles (or abscission site) was measured manually using a calliper.

To quantify plant height, primary shoots of 20 plants from each of the three lines were measured from soil level to growth tip using a flexible tape measure to account for plant curvature.

To quantify fecundity, developing fruits (those with inflating calyces), open flowers, and closed flower buds we counted across all shoots on 15 plants from each of the lines.

To quantify fruit weight, mature fruits (yellow/orange in color) were weighed without calyx. Fruits were weighed in cluster groups of five fruits for ten plant replicates (for a total of 50 fruits). Cluster weight was divided by five to calculate average fruit weight per plant.

**Statistical analysis**

Statistical analyses of the phenotypes were performed using t-test. Significance was determined as p-value $< 0.05$.
